# Reward motivation increases univariate activity but has limited effect on coding of task-relevant information across the frontoparietal cortex

**DOI:** 10.1101/609537

**Authors:** Sneha Shashidhara, Yaara Erez

## Abstract

Selection and integration of information based on current goals is fundamental for goal-directed behavior. Reward motivation has been shown to improve behavioral performance, yet the neural mechanisms that link motivation and control processes, and in particular its effect on context-dependent information processing, remain unclear. We used functional magnetic resonance imaging (fMRI) in 24 human volunteers (13 females) to test whether reward motivation enhances the coding of task-relevant information across the frontoparietal cortex, as would be predicted based on previous experimental evidence and theoretical accounts. In a cued target detection task, participants detected whether an object from a cued visual category was present in a subsequent display. The combination of the cue and the object visual category determined the behavioral status of the objects. To manipulate reward motivation, half of all trials offered the possibility of a monetary reward. We observed an increase with reward in overall univariate activity across the frontoparietal control network when the cue and subsequent object were presented. Multivariate pattern analysis (MVPA) showed that behavioral status information for the objects was conveyed across the network. However, in contrast to our prediction, reward did not increase the discrimination between behavioral status conditions in the stimulus epoch of a trial when object information was processed depending on a current context. In the high-level general-object visual region, the lateral occipital complex, the representation of behavioral status was driven by visual differences and was not modulated by reward. Our study provides useful evidence for the limited effects of reward motivation on task-related neural representations and highlights the necessity to unravel the diverse forms and extent of these effects.

## 1. Introduction

A fundamental aspect of flexible goal-directed behavior is the selection and integration of information depending on a current goal to determine its relevance to behavior and lead to a decision. With growing interest in recent years in the link between cognitive control and reward motivation, it has been proposed that reward enhances control processes by sharpening representation of task goals and prioritizing task-relevant information across the frontoparietal control network and other regions associated with cognitive control (Botvinick and Braver, 2015; Etzel et al., 2016; Kruglanski et al., 2002; Simon, 1967). In line with this idea, it has been shown that motivation, usually manipulated as monetary reward, increases task performance (Padmala and Pessoa, 2011, 2010). Neuroimaging studies linked increased activity with reward in frontoparietal regions across a range of tasks, including working memory (Pochon et al., 2002; Taylor et al., 2004), selective attention (Krebs et al., 2012; Mohanty et al., 2008), response inhibition (Padmala and Pessoa, 2011), and problem solving (Shashidhara et al., 2019).

Although the accumulating evidence at the behavioral and neural level in humans are consistent with this sharpening and prioritization account (Braver, 2012; Chiew and Braver, 2014; Kruglanski et al., 2002; Miller et al., 1960; Pessoa, 2009; Simon, 1967), they do not directly address the effect of reward motivation on the coding of task-related information and selection and integration processes. Some support for this idea comes from single-neuron data recorded from the prefrontal cortex of non-human primates: reward was associated with greater spatial selectivity, enhanced activity related to working memory and modulated task-related activity based on the type of reward (Kennerley and Wallis, 2009; Leon and Shadlen, 1999; Watanabe, 1996). A more direct evidence in humans was recently demonstrated by Etzel et al. (2016). They showed that reward enhances coding of task cues across the frontoparietal cortex and suggested that task-set efficacy increases with reward. It remained an open question whether this facilitative effect of reward is limited to preparatory cues, or if reward also enhances the coding of behaviorally relevant information when the cue and a subsequent stimulus are integrated, leading to the behavioral decision. Effects of reward during both cue and stimulus epochs of a trial are complementary to one another, and processing during both epochs is vital when reaching a decision. In a recent electroencephalogram (EEG) study, Hall-McMaster et al. (Hall-McMaster et al., 2019) showed that reward increases coding of task cues. However, these increases were observed only for ‘switch’ trials where task rules had to be updated. In addition, they also provided some evidence that the representation of task-relevant features is enhanced when reward level is high. Given the limited spatial resolution of EEG, it is unclear whether these effects of reward are specific to the frontoparietal control network. Taken together, these previous findings demonstrate that the effects of reward on representation of task-related information may not be generalized and robust but rather specific to certain cognitive demands and trial epochs.

To further shed light on the extent of the effects of reward on neural representations, here we focus on the attentional saliency of task-relevant information as determined by the nature of the task. Consistent with the idea that the frontoparietal network is involved in selection and integration of task-relevant information, in previous work we showed that top-down attentional saliency of items from different visual categories, as determined by their likelihood of being targets, drives representation across this network (Erez and Duncan, 2015). Thus, rather than representation of the visual categories themselves, it was the attentional saliency, or behavioral status of the items, as determined by the integration of cue and stimulus input, that was the task-relevant aspect represented across the network. In the current study, we ask whether reward motivation enhances the representation of task-related behavioral status across the frontoparietal network. Furthermore, previous studies have associated reward with decreased conflict in interference tasks (Krebs et al., 2013; Padmala and Pessoa, 2011; Stürmer et al., 2011), suggesting that any effect of reward may be particularly important for high-conflict items, in other words, a conflict-contingent effect. We therefore also asked whether such facilitative effect of reward is selective for highly conflicting items. We used fMRI and multivariate pattern analysis (MVPA) to measure representation of the task-related behavioral status as well as visual categories in distributed patterns of response in the human brain. Specifically, we tested whether pattern discriminability increased with reward in the *a priori* chosen frontoparietal ‘multiple-demand’ (MD) network (Duncan, 2010; Fedorenko et al., 2013), which has been associated with multiple aspects of cognitive control, such as working memory, task sets, conflict monitoring, task switching, and task-dependent categorical decisions (Cole et al., 2016; Erez and Duncan, 2015; Fedorenko et al., 2013; Li et al., 2007; Muhle-Karbe et al., 2017; Nastase et al., 2017; Vergauwe et al., 2015; Wisniewski et al., 2016). Lastly, it is commonly accepted that top-down signals from the frontoparietal MD network to the visual cortex play an important role in the processing of task-related information. Therefore, we tested whether similar effects of reward would be observed in the high-level general-object visual region, the lateral occipital complex (LOC).

## 2. Materials and Methods

### 2.1 Participants

24 participants (13 females), between the ages of 18-40 years (mean age: 25) took part in the study. Four additional participants were excluded due to large head movements during the scan (greater than 5 mm). The sample size was determined prior to data collection as typical for neuroimaging studies, in accordance with sample sizes in previous fMRI studies that showed sufficient power to detect representation of behavioral status and effects of reward on coding of task information (Erez and Duncan, 2015; Etzel et al., 2016), and to comply with counter-balancing requirements of the experimental design across participants. All participants were right-handed with normal or corrected-to-normal vision and had no history of neurological or psychiatric illness. The study was conducted with approval by the Cambridge Psychology Research Ethics Committee. All participants gave written informed consent and were monetarily reimbursed for their time.

### 2.2 Task design

Participants performed a cued target detection categorization task in the MRI scanner (Figure 1A). On each trial, integration of a preceding cue and the presented object determined the behavioral status of the object. Our primary question concerned the representation during the stimulus epoch of a trial where cue and stimulus are integrated, and we therefore designed the task accordingly. At the beginning of each trial, one of three visual categories (sofas, shoes, cars) was cued, determining the target category for that trial. Participants had to indicate whether the subsequent object matched this category or not by pressing a button. For each participant, only two of the categories were cued as targets throughout the experiment. Depending on the cue on a given trial, objects from these categories could be either Targets, or nontargets with high conflict as they could serve as targets on other trials (High-conflict nontarget). The third category was never cued, therefore objects from this category served as Low-conflict nontargets. This design yielded three behavioral status conditions: Targets, High-conflict nontargets and Low-conflict nontargets (Figure 1B). Critically, following the integration of the cue and presented object, the relevant information that is expected to be represented across the MD network is the behavioral status of a given category, rather than the visual category itself (Erez and Duncan, 2015). Therefore, the task-relevant distinctions that were tested using MVPA were pairs of categories with different behavioral status. The assignment of the categories to be cued (and therefore serve as either Targets or High-conflict nontargets) or not (and serve as Low-conflict nontargets) was counter-balanced across participants.

**Figure 1:**
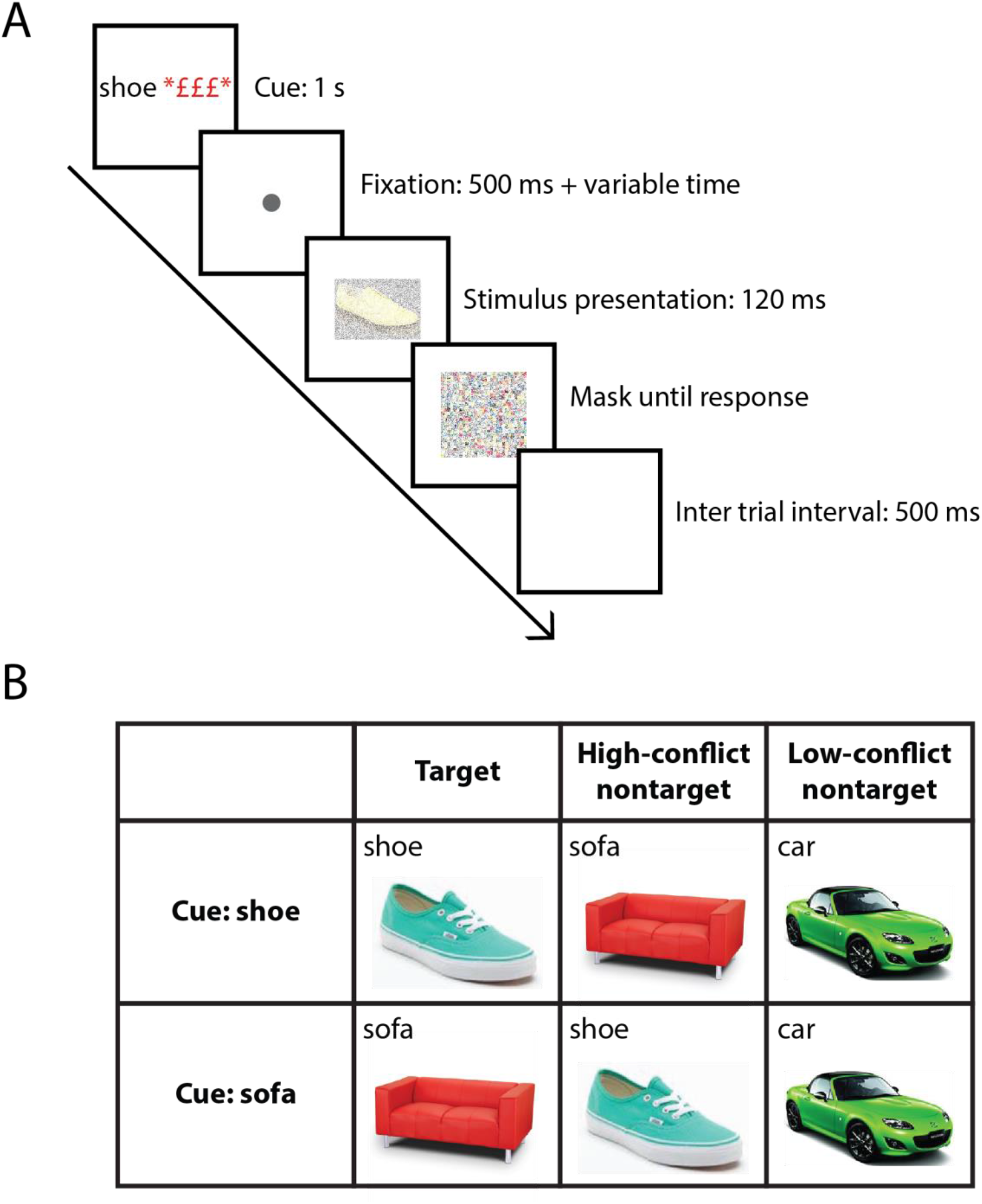
Experimental paradigm. **A.** An example trial. A trial began with a cue (1 s) indicating the target category, followed by 500 ms fixation period. Reward trials were cued with three red £ symbols next to the target category. After an additional variable time (0.4, 0.7, 1.0 or 1.3 s), an object was presented for 120 ms. The object was then masked (a scramble of the all the stimuli used), until response or for a maximum of 3 s. The participants pressed a button to indicate whether the object was from the cued category (Target trials) or not (Nontarget trials). 33% of all trials were catch trials. These included cue followed by fixation dot for 500 ms, which then turned red for another 500 ms indicating the absence of the stimulus, followed by the inter-trial interval. **B.** Experimental conditions. For each participant, two categories served as potential targets depending on the cue, and a third category never served as target. Here as an example, shoes and sofas are the cued categories and cars as the uncued category. In the Target trials, the presented object matched the cued category. In the High-conflict nontarget trials, the object did not match the cued category, but was from the other cued category, therefore could serve as a target on other trials. In the Low-conflict nontarget trials, the presented object was from the category that was never cued. Overall, this design yielded three levels of behavioral status: Targets, High-conflict nontargets, and Low-conflict nontargets. The design was used for both no-reward and reward conditions.

To manipulate motivation, half of the trials were cued as reward trials in which a monetary reward was offered. On these reward trials, participants had the chance of earning £1 if they completed the trial correctly and within a time limit. Four random reward trials out of 32 in each run were assigned the £1 reward, thus ensuring an incentive on all reward trials. To avoid longer reaction times when participants try to maximize their reward, a response time threshold was used for reward trials, set separately for each participant as the average of 32 trials in a pre-scan session. The participants were told that the maximum reward they could earn is £24 in the entire session (£4 per run) and were not told what the time threshold was. Therefore, to maximize their gain, participants had to treat every reward trial as a £1 trial and respond as quickly and as accurately as possible, just as in no-reward trials.

Each trial started with a 1 s cue, which was the name of a visual category that served as the target category for this trial. On reward trials, the cue included three red pound signs presented next to the category name. The cue was followed by a fixation dot in the center of the screen presented for 0.5 s and an additional variable time of either 0.1, 0.4, 0.7 or 1 s, selected randomly, in order to make the stimulus onset time less predictable. The stimulus was then presented for 120 ms and was followed by a mask. The stimulus duration was fixed and identical for all trials. Participants indicated by a button press whether this object belonged to the cued target category (present) or not (absent). Following response, a 1 s blank inter-trial interval separated two trials. For both reward and no-reward trials, response time was limited to a maximum of 3 s, after which the 1 s blank inter-trial interval started even when no response was made. For reward trials, an additional subject-specific response time threshold was used as mentioned above to determine whether the participants earned the reward or not, but this time threshold did not affect the task structure and was invisible to the participants.

We used catch trials to decorrelate the BOLD signals of the cue and stimulus phases. 33% of all trials included cue followed by fixation dot for 500 ms, which then turned red for another 500 ms indicating the absence of the stimulus, followed by the inter-trial interval.

The combination of multiple factors in the task design resulted in a cognitively demanding task that was expected to recruit MD regions. These factors include frequent cue changes, short stimulus presentation duration, short inter-trial interval and change of response mapping. We also note that there are multiple aspects of information selection that are of interest when considering the effect of reward motivation on neural representation, one of which is how it may bias attention to reward-associated stimuli when two or more stimuli are presented simultaneously and compete for attention. Here we focus on single stimulus displays and the integration of cue and stimulus that generates the behavioral status of the stimuli as a starting point, while attentional competition may be address in future studies.

### 2.3 Stimuli

Objects were presented at the center of the screen on a grey background. The objects were 2.95° visual angle along the width and 2.98° visual angle along the height. Four exemplars from each visual category were used. Exemplars were chosen with similar colors, dimensions, and orientation across the categories. All exemplars were used an equal number of times in each condition and in each run to ensure that any differences between the experimental conditions will not be driven by the variability of exemplars. To increase the task demand, based on pilot data, we added Gaussian white noise to the stimuli. The post-stimulus mask was generated by randomly combining pieces of the stimuli that were used in the experiment. The mask was the same size as the stimuli and was presented until a response was made or the response time expired.

### 2.4 Structure and design

Each participant completed 6 functional runs of the task in the scanner (mean duration ± SD: 6.2 ± 0.13 min). Each run started with a response-mapping instructions screen (e.g., left = target present, right = target absent), displayed until the participants pressed a button to continue. Halfway through the run, the instructions screen was presented again with the reversed response mapping. All trials required a button response to indicate whether the target was present or absent, and the change of response mapping ensured that conditions were not confounded by the side of the button press. Each run included 104 trials. Out of these, 8 were dummy trials following the response mapping instructions (4 after each instructions screen) and were excluded from the analysis. Of the remaining 96 trials, one-third (32 trials) were cue-only trials (catch trials). Of the remaining 64 trials, 32 were no-reward trials and 32 were reward trials. Of the 32 no-reward trials, half (16) were cued with one visual category, and half (16) with the other. For each cued category, half of the trials (8) were Target trials, and half of the trials (8) were nontarget trials, to ensure an equal number of target (present) and nontarget (absent) trials. Of the nontarget trials, half (4) were High-conflict nontargets, and half (4) were Low-conflict nontargets. There were 4 trials per cue and reward level for the High- and Low-conflict nontarget conditions, and 8 for the Target condition, with the latter split into two regressors (see 2.7.2 General Linear Model (GLM) for the Main Task section below). A similar split was used for reward trials. The task included an overall of 96 Target trials, 48 High-conflict nontarget trials, and 48 Low-conflict nontarget trials for each reward level (no-reward and reward) across runs and cued categories. An event-related design was used, and the order of the trials was randomized in each run. At the end of each run, the money earned in the reward trials and the number of correct trials (across both reward and no-reward trials) were presented on the screen.

### 2.5 Functional localizers

In addition to the main task, we used two other tasks in order to functionally localize MD regions and LOC in individual participants using independent data. These functional localizer data were used for subject-specific ROI definitions within anatomical constraints of ROI templates for MD regions and LOC (Fedorenko et al., 2010; Shashidhara et al., 2020). See 2.7.5 ROI definition and 2.7.6 Subject-specific ROIs for more details. For consistency, this approach was used for all the univariate and multivariate pattern analyses.

To localize MD regions, we used a spatial working memory task (Fedorenko et al., 2013). This task has been previously shown to robustly recruit the MD network similarly to other tasks across multiple cognitive domains (Fedorenko et al., 2013). Furthermore, when used as a localizer task in conjunction with anatomical constraints (i.e., the same approach as we use in the current study), decoding levels in an independent task across the MD network were similar to other MD localizer tasks (Shashidhara et al., 2020), thus supporting its suitability for subject-specific localization of the MD network, in particular when addressing pattern discriminability. On each trial, participants remembered 4 locations (Easy condition) or 8 locations (Hard condition) in a 3X4 grid. Each trial started with fixation for 500 ms. Locations on the grid were then highlighted consecutively for 1 s (1 or 2 locations at a time, for the Easy and Hard conditions, respectively). In a subsequent two-alternative forced-choice display (3 s), participants had to choose the grid with the correct highlighted locations by pressing the left or the right button. Feedback was given after every trial for 250 ms. Each trial was 8 s long, and each block included 4 trials (32 s). There was an equal number of correct grids on the right and left in the choice display. Participants completed 2 functional runs of 5 min 20 sec each, with 5 Easy blocks alternated with 5 Hard blocks in each run. We used the contrast of Hard vs. Easy blocks to localize MD regions.

As a localizer for LOC, we used a one-back task with blocks of objects interleaved with blocks of scrambled objects. The objects were in grey scale and taken from a set of 61 everyday objects (e.g., camera, coffee cup, etc.). Participants had to press a button when the same image was presented twice in a row. Images were presented for 300 ms followed by a 500 ms fixation. Each block included 15 images with two image repetitions and was 12 s long. Participants completed two runs of this task, with 8 object blocks, 8 scrambled object blocks, and 5 fixation blocks. The objects vs. scrambled objects contrast was used to localize LOC.

### 2.6 Scanning session

The scanning session included a structural scan, 6 functional runs of the main task, and 4 functional localizer runs – 2 for MD regions and 2 for LOC. The scanning session lasted up to 100 minutes, with an average 65 minutes of EPI time. The tasks were introduced to the participants in a pre-scan training session. The average reaction time of 32 no-reward trials of the main task completed in this practice session was set as the time threshold for the reward trials to be used in the scanner session. All tasks were written and presented using Psychtoolbox3 (Brainard, 1997) and MatLab (The MathWorks, Inc).

#### 2.6.1 Data acquisition

fMRI data were acquired using a Siemens 3T Prisma scanner with a 32-channel head coil. We used a multi-band imaging sequence (CMRR, release 016a) with a multi-band factor of 3, acquiring 2 mm isotropic voxels (Feinberg et al., 2010). Whole-brain scans were acquired. Other acquisition parameters were: TR = 1.1 s, TE = 30 ms, 48 slices per volume with a slice thickness of 2 mm and no gap between slices, in plane resolution 2 × 2 mm, field of view 205 mm, flip angle 62°, and interleaved slice acquisition order. No iPAT or in-plane acceleration were used. T1-weighted multiecho MPRAGE (van der Kouwe et al., 2008) high-resolution images were also acquired for all participants, in which four different TEs were used to generate four images (voxel size 1 mm isotropic, field of view of 256 × 256 × 192 mm, TR = 2530 ms, TE = 1.64, 3.5, 5.36, and 7.22 ms). The voxelwise root mean square across the four MPRAGE images was computed to obtain a single structural image.

### 2.7 Data and statistical analysis

The primary analysis approach was multi-voxel pattern analysis (MVPA), to assess representation of behavioral status distinctions with and without reward. An additional ROI-based univariate analysis was conducted to confirm the recruitment of the MD network. Preprocessing, GLM and univariate analysis of the fMRI data were performed using SPM12 (Wellcome Department of Imaging Neuroscience, London, England; www.fil.ion. ucl.ac.uk), and the Automatic Analysis (aa) toolbox (Cusack et al., 2014). The analysis choices described below were determined *a priori* to conducting the analyses.

We used an alpha level of .05 for all statistical tests. Bonferroni correction for multiple comparisons was used when required, and the corrected p-values and uncorrected t-values are reported. All t tests that were used to compare two conditions were paired due to the within-subject design. A one-tailed t test was used when the prediction was directional, including testing for classification accuracy above chance level. All other t tests in which the *a priori* hypothesis was not directional were two-tailed. Additionally, effect size (Cohen’s *d_z_*) was computed. We note that using a t test for group level inference of classification accuracy above chance may be limited in its interpretation as true accuracies cannot be below chance level (Allefeld et al., 2016). We emphasize that tests for classification accuracies above chance are reported in the Results as a complementary measure while the main research question concerns the *change* in decoding level with reward. A post-hoc sensitivity power analysis was conducted for the effect of reward modulation of decoding in MD regions, with alpha level of 0.05, power of 0.8 and one tail (Faul et al., 2007). All analyses were conducted using custom-made MATLAB (The Mathworks, Inc) scripts, unless otherwise stated.

#### 2.7.1 Pre-processing

Initial processing included motion correction and slice time correction. The structural image was coregistered to the Montreal Neurological Institute (MNI) template, and then the mean EPI was coregistered to the structural. The structural image was then normalized to the MNI template via a nonlinear deformation, and the resulting transformation was applied on the EPI volumes. Spatial smoothing of FWHM = 5 mm was performed for the functional localizers data only.

#### 2.7.2 General Linear Model (GLM) for the main task

We used GLM to model the main task and localizers’ data. Regressors for the main task included 12 conditions during the stimulus epoch and 4 conditions during the cue epoch. Regressors during the stimulus epoch were split according to reward level (no-reward, reward), cued visual category (cue 1, cue 2), and behavioral status (Target, High-conflict nontarget, Low-conflict nontarget). Since the combination of cued category and the stimulus category on each trial determined the behavioral status, the regressors for each reward level included all possible combinations of cued and stimulus category: cue1-stimulus1 (Target), cue1-stimulus2 (High-conflict nontarget), cue1-stimulus3 (Low-conflict nontarget), cue2-stimulus 2 (Target), cue2-stimulus1 (High-conflict nontarget), cue2-stimulus2 (Low-conflict nontarget). To ensure an equal number of target present and target absent trials, the number of Target trials in our design was twice the number of High-conflict and Low-conflict nontarget trials. The Target trials included two repetitions of each combination of cue, visual category and exemplar, with a similar split for reward trials. These two Target repetitions were modelled as separate Target1 and Target2 regressors in the GLM to make sure that all the regressors were based on an equal number of trials but were invisible to the participants. All the univariate and multivariate analyses were carried out while keeping the two Target regressors separate to avoid any bias of the results, and they were averaged at the final stage of the results. Overall, the GLM included 16 regressors of interest for the 12 stimulus conditions. Each regressor was based on data from all correct trials in the respective condition in each run (up to 4 trials), i.e., the regressors were computed separately for each run. To account for possible effects of reaction time (RT) on the beta estimates because of the varying duration of the stimulus epoch, and as a consequence their potential effect on decoding results, these regressors were modelled with durations from stimulus onset to response (Woolgar et al., 2014). This model assumes that activity in each voxel continues throughout the phase, and as long as the participant is performing the trial. This ensured that we capture the entire cognitive processing of integration of stimulus and cue until a decision is reached in all trials, but we emphasize that the actual stimulus duration was fixed for all trials. Since this model scales the regressors based on the reaction time, the beta estimates reflect activation per unit time and are comparable across conditions with different durations. In addition to regressors during the stimulus epoch, we also modelled the cue epoch. Cue regressors included both task and cue-only (catch) trials and were split by reward level and cued category, modelled with duration of 1 s. Cue regressors were based on 16 trials per regressor per run. As one-third of all trials were catch trials, the cue and stimulus epoch regressors were decorrelated and separable in the GLM. All the regressors were convolved with the canonical hemodynamic response function (HRF). The 6 movement parameters and run means were included as covariates of no interest.

As one of our control analyses, we classified patterns of activity of motor response (left vs. button presses) in motor areas. For this analysis, we constructed a separate GLM. This GLM was similar to the above GLM of the main task, with the only difference being the stimulus phase regressors. For this model, we used two regressors (right press, left press) for each of the two reward levels (no-reward, reward), a total of four regressors.

#### 2.7.3 GLM for the functional localizers

For the MD localizer, regressors included Easy and Hard blocks. For LOC, regressors included objects and scrambled objects blocks. Each block was modelled with its duration. The regressors were convolved with the canonical hemodynamic response function (HRF). The 6 movement parameters and run means were included as covariates of no interest.

#### 2.7.4 Univariate analysis

We conducted an ROI analysis to test for the effect of reward on overall activity for the different behavioral status conditions and cues. We used individually defined ROIs based on subject-specific independent functional localizer data within anatomical constraints of templates for the MD network and for LOC as defined below (see 2.7.5 ROI definition 2.7.6 and subject-specific ROIs). Using the MarsBaR toolbox (http://marsbar.sourceforge.net; Brett et al., 2002) for SPM 12, beta estimates for each regressor of interest were extracted and averaged across runs, and across voxels within each ROI, separately for each participant and condition. For the MD network, beta estimates were also averaged across hemispheres (see ROI definition below). Second-level analysis was done on beta estimates across participants using repeated measures ANOVA. The data for the Target condition was averaged across the two Target1 and Target2 regressors, separately for the no-reward and reward conditions.

#### 2.7.5 ROI definition

##### MD network template

ROIs of the MD network were defined *a priori* using an independent data set (Fedorenko et al. 2013; see t-map at http://imaging.mrc-cbu.cam.ac.uk/imaging/MDsystem). These included the anterior, middle, and posterior parts of the middle frontal gyrus (aMFG, mMFG, and pMFG, respectively), a posterior dorsal region of the lateral frontal cortex (pdLFC), AI-FO, pre-SMA/ACC, and IPS, defined in the left and right hemispheres. The visual component in this template is widely accepted as a by-product of using largely visual tasks and is not normally considered as part of the MD network. Therefore, it was not included in the analysis. The MD network is highly bilateral, with similar responses in both hemispheres (Fedorenko et al., 2013). We therefore averaged the results across hemispheres in all the analyses.

##### LOC template

LOC was defined using data from a functional localizer in an independent study with 15 participants (Lorina Naci, PhD dissertation, University of Cambridge). In this localizer, forward- and backward-masked objects were presented, as well as masks alone. Masked objects were contrasted with masks alone to identify object-selective cortex (Malach et al., 1995). Division to the anterior part of LOC, the posterior fusiform region (pFs) of the inferior temporal cortex, and its posterior part, the lateral occipital region (LO) was done using a cut-off MNI coordinate of Y=-62, as previous studies have shown differences in processing for these two regions (Erez and Yovel, 2014; MacEvoy and Epstein, 2011).

##### Motor regions template

For one of our control analyses, we used two anatomical masks for motor regions from the FSL (FMRIB Software Library) Harvard-Oxford atlas (Desikan et al., 2006).The precentral gyrus and the supplementary motor cortex.

#### 2.7.6 Subject-specific ROIs

To define subject-specific ROIs, we used independent functional localizer data for each subject within the anatomical constraints of the MD network and LOC templates (Fedorenko et al., 2010; Shashidhara et al., 2020). This has allowed us to use both a template, consistent across participants, as well as subject-specific data as derived from the functional localizers. This approach was used for all the univariate and multivariate pattern analyses in the MD network and the LOC. For the multivariate analysis, using this approach also ensured controlled ROI size for comparison between regions within the MD network and between sub-regions in LOC. For each participant, beta estimates of each condition and run were extracted for each ROI based on the MD network and LOC templates. For each MD ROI, we then selected the 200 voxels with the largest t-value for the Hard vs. Easy contrast as derived from the independent subject-specific functional localizer data. This number of voxels was defined prior to any data analysis (Erez and Duncan, 2015; Shashidhara et al., 2020). The 200 voxels captured 19.5% of voxels in each MD ROI on average. For each LOC sub-region, we selected 180 voxels with the largest t-values of the object vs. scrambled contrast from the independent subject-specific functional localizer data. The selected voxels were used for the voxelwise patterns in the MVPA for the main task. The number of voxels that was used for LOC was smaller than for MD regions because of the size of the pFs and LO masks. For the analysis that addressed decoding of visual categories and compared MD regions with the visual regions, we used 180 voxels from all regions to keep the ROI size the same. On average, the 180 voxels captured 17.5% of voxels in each MD ROI and 45% of each LOC ROI. Figure S1 shows probabilities maps of the distribution of subject-specific ROIs across MD and LOC ROIs.

To ensure that the decoding results are robust and do not depend on the choice of the number of selected voxels in each ROI, we repeated the main decoding analysis for MD regions with a range of ROI sizes (100, 150, 250 and 300 voxels).

#### 2.7.7 ROI-based multivoxel pattern analysis (MVPA)

We used MVPA to test for the effect of reward motivation on pattern discriminability between the task-related behavioral status pairs. Voxelwise patterns using the selected voxels within each template were computed for all the task conditions in the main task. We applied our classification procedure on all possible pairs of conditions as defined by the GLM regressors of interest during the stimulus presentation epoch, for the no-reward and reward conditions separately (Figure 1B). The beta estimates for each condition and run were used for the MVPA and were not normalized at any point during the analysis process. For each pair of conditions, MVPA was performed using a support vector machine classifier (LIBSVM library for MATLAB, c=1) implemented in the Decoding Toolbox (Hebart et al., 2015). We used leave-one-run-out cross-validation in which the classifier was trained on the data of five runs (training set) and tested on the sixth run (test set). This was repeated 6 times, leaving a different run to test each time, and classification accuracies were averaged across these 6 folds. Classification accuracies were then averaged across pairs of different cued categories, yielding discrimination measures for three pairs of behavioral status (Targets vs. High-conflict nontargets, Targets vs. Low-conflict nontargets, and High-conflict vs. Low-conflict nontargets) within each reward level (no-reward, reward). Because the number of Target trials in our design was twice the number of High-conflict and Low-conflict nontarget trials, each discrimination that involved a Target condition was computed separately for the two Target regressors (Target1 and Target2) and classification accuracies were averaged across them. We note that for each participant, while the visual categories were balanced for Targets and High-conflict nontargets, the visual category that was assigned as Low-conflict nontarget did not serve as a Target or High-conflict nontarget. Nevertheless, based on previous work, it is expected that behavioral status will be represented across the MD network with little or no effect of the visual category itself (Erez and Duncan, 2015).

The Target and High-conflict nontarget pairs of conditions included cases when both conditions had an item from the same visual category as the stimulus (following different cues), as well as cases in which items from two different visual categories were displayed as stimuli (following the same cue). To test for the contribution of the visual category to the discrimination, we split the Target vs. High-conflict nontarget pairs of conditions into these two cases and the applied statistical tests accordingly.

Although our main hypothesis addressed the behavioral status of the presented stimuli, several studies showed that information related to stimulus features is also represented across the MD network (Jackson et al., 2017; Woolgar et al., 2015). Rather than increased representation of the behavioral status with reward as we hypothesized, information about the visual categories themselves could increase. We therefore also tested for decoding of visual categories when both stimuli are Targets or both are High-conflict nontargets, and whether this is modulated by reward.

To ensure that our results for the main study question are robust across different pattern analysis techniques, we tested for the modulation of the representation of behavioral status by reward using a linear discriminant contrast (LDC) (Carlin and Kriegeskorte, 2017; Nili et al., 2014), in addition to SVM. The LDC is the cross-validated Mahalanobis distance between two classes and provides unbiased distances. Larger LDC indicates greater pattern dissimilarity, i.e., greater discriminability, and 0 reflects no discriminability. The MVPA analysis as described above was repeated with the LDC measure and using the same voxel selection procedure. For each participant, LDC was computed for all possible pairs of conditions and LDC values were then averaged as described above to address each of the questions. The LDC analysis was implemented using the Representational Similarity Analysis toolbox (Nili et al., 2014).

As one of our control analyses, we tested for classification of right vs. left button presses in motor regions for no-reward and reward trials. This analysis was used as a quality check for the data and the correctness of the MVPA pipeline. We used beta estimates from a separate GLM in which trials were modelled for right/left button press only for each of the two reward levels. Decoding accuracies were computed for each reward level separately and then averaged across reward levels. The analysis was performed similarly to the MVPA described above using a support vector machine classifier (LIBSVM library for MATLAB, c=1), implemented in the Decoding Toolbox (Hebart et al., 2015), and using a leave-one-run-out cross-validation. All voxels within the motor ROIs were used.

#### 2.7.8 Whole-brain searchlight pattern analysis

To test whether regions outside the MD network show change in discriminability between voxelwise patterns of activity of task-related behavioral status when reward is introduced, we conducted a whole-brain searchlight pattern analysis (Kriegeskorte et al., 2006) using the Decoding Toolbox (Hebart et al., 2015). This analysis enables the identification of focal regions that carry relevant information, unlike the ROI-based analysis, which tests for a more widely distributed representation of information across larger ROIs. For each participant, data was extracted from spherical ROIs with a 5 mm radius (maximum 65 voxels), centered on each voxel in the brain. These voxels were used to perform the same MVPA analysis using SVM as described above. Thus, for each voxel, we computed the classification accuracies for the relevant behavioral status distinctions, separately for the reward and no-reward conditions. These whole-brain maps were smoothed using a 5 mm FWHM Gaussian kernel. The t-statistic from a second level random-effects analysis on the smoothed maps was thresholded at the voxel level using FDR correction (p < 0.05).

#### 2.7.9 Data and code accessibility

All raw data and code used in this study will be deposited on a departmental repository upon publication and will be available upon any request.

## 3. Results

### 3.1 Behavior

Overall accuracy levels were high (mean ± SD: 92.51% ± 0.08%). Mean and SD accuracy rates for the Target, High-conflict nontarget and Low-conflict nontarget conditions in the no-reward trials were 91.2% ± 5.8%, 89.1% ± 8.8%, and 96.6% ± 3.8%, respectively; and for the reward trials they were 94.2% ± 5.0%, 87.8%± 8.7%, 96.1% ± 4.4%, respectively (Figure 2A). A two-way repeated measures ANOVA with reward level and behavioral status as within-subject factors showed no main effect of reward (*F*_1, 23_ = 0.49, *p* = 0.49), confirming that the added time constraint for reward trials did not lead to drop in performance. There was a main effect of behavioral status (*F*_2, 23_ = 29.64, *p* < 0.001) and an interaction between reward level and behavioral status (*F*_2, 23_ = 5.81, *p* < 0.01). Post-hoc tests with Bonferroni correction for multiple comparisons showed larger accuracies for Low-conflict nontargets compared to Targets and High-conflict nontargets in the no-reward trials (Two-tailed t-test: *t_23_* = 5.64, *p* < 0.001, *d_z_* = 1.15; *t_23_* = 5.50, *p* < 0.001, *d_z_* = 1.12 respectively), as expected given that the Low-conflict nontarget category was fixed throughout the experiment. In the reward trials, performance accuracies were larger for Targets compared to High-conflict nontargets (*t_23_* = 4.45, *p* < 0.001, *d_z_* = 0.91) and Low-conflict nontargets compared to High-conflict ones (*t_23_* = 5.92, *p* < 0.001, *d_z_* = 1.2), with only marginal difference between Targets and Low-conflict nontargets (*t_23_* = 2.49, *p* = 0.06). Accuracies for Target trials were larger for the reward trials compared to no-reward (*t_23_* = 2.92, *p* = 0.008, *d_z_* = 0.61), indicating a possible behavioral benefit of reward. There was no difference between reward and no-reward trials for High-conflict and Low-conflict nontargets (*t_23_* < 1.1, *p* > 0.1, for both).

**Figure 2:**
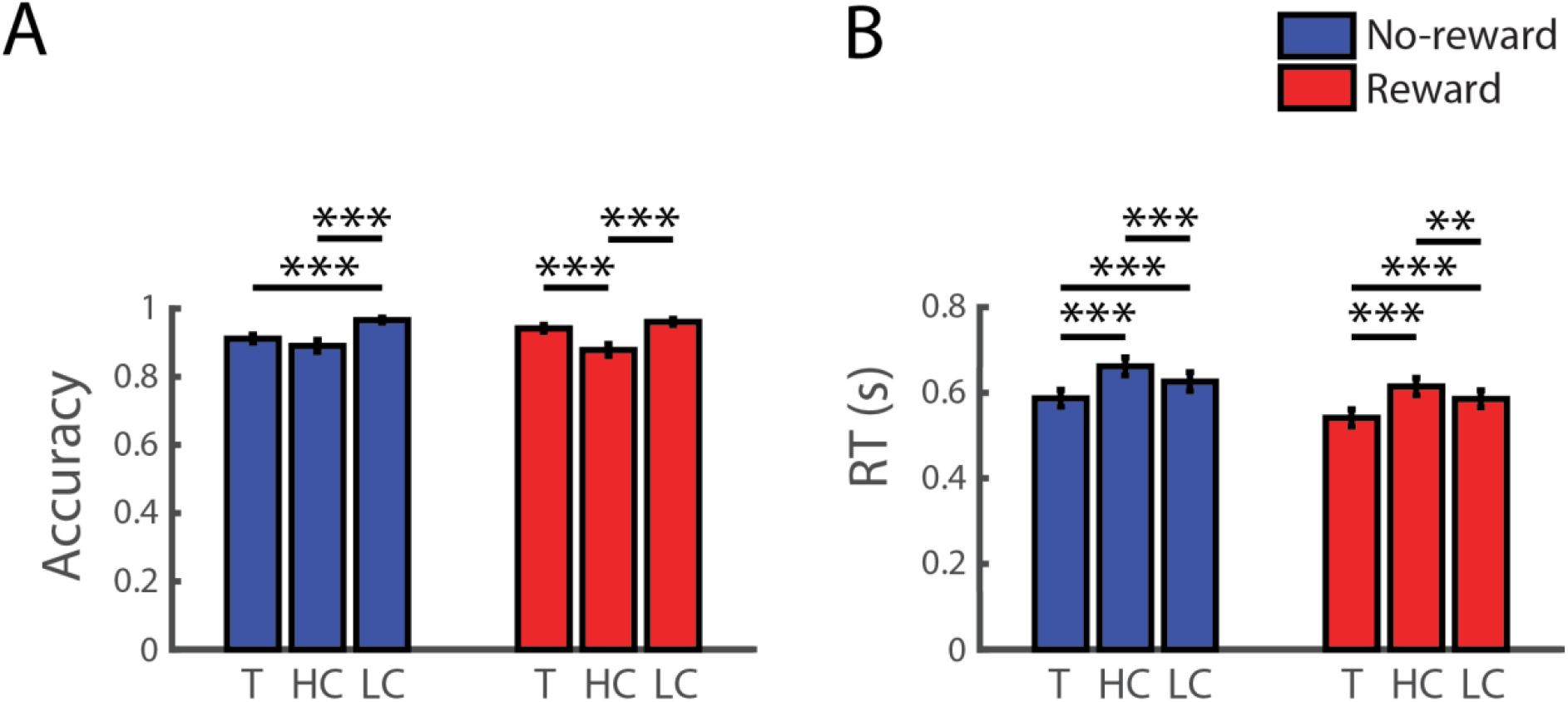
Behavioral results. **A.** Accuracy rates. Proportion of correct trials is presented for Targets (T), High-conflict nontargets (HC) and Low-conflict nontargets (LC) for no-reward and reward trials. **B.** Reaction times. Error bars indicate S.E.M. Asterisks show significance levels following a two-tailed paired t-test, corrected for 3 comparisons. ** *p* < 0.01, *** *p* < 0.001.

RT of successful trials for the three behavioral status conditions, Target, High-conflict nontarget and Low-conflict nontarget, in the no-reward trials were 589 ± 98 ms, 662 ± 103 ms, and 626 ± 107 ms, respectively (mean ± SD); RTs for these conditions in the reward trials were 541 ± 99 ms, 614 ± 99 ms, 585 ± 97 ms, respectively (mean ± SD) (Figure 2B). A two-way repeated measures ANOVA with reward level (no-reward, reward) and behavioral status as within-subject factors showed a main effect of reward (*F*_1, 23_ = 40.07, *p* < 0.001), with reward trials being shorter than no-reward trials, as expected from the experimental design that required a response within a time limit to receive the reward. An additional main effect of behavioral status (*F*_2, 23_ = 50.97, *p* < 0.001) was observed, with no interaction between reward and behavioral status (*F*_2, 23_ = 0.63, *p* = 0.54). Subsequent post-hoc tests with Bonferroni correction for multiple comparisons showed that RTs for Target trials were faster than High-conflict and Low-conflict nontarget trials (*t_23_* = 10.03, *p* < 0.001, *d_z_* = 2.05; *t_23_* = 5.17, *p* < 0.001, *d_z_* = 1.06 respectively), and Low-conflict nontarget trials were faster than the High-conflict ones (*t_23_* = 4.96, *p* < 0.001, *d_z_* = 1.01), as expected from a cued target detection task.

### 3.2 Activity across the MD network during the cue epoch

To address our primary research question, the analysis focused on the stimulus epoch. However, to get a full picture of the data and for comparability with previous studies that showed increase in cue information, we also report the results for the cue epoch here. This analysis focuses only on the MD network and not the LOC, since no object stimuli were presented at this epoch of the trial.

We first tested for a univariate effect of reward during the cue phase (averaged across the β estimates of the two cues) across the subject-specific ROIs for all MD regions. A two-way repeated measures ANOVA with reward (2: no-reward, reward) and ROI (7) as factors showed a main effect of reward (*F*_1, 23_ = 20.53, *p* < 0.001) with increased activity during the reward trials compared to the no-reward trials. There was also a main effect of ROI (*F*_6, 138_ = 11.63, *p* < 0.001) and an interaction of reward and ROI (*F*_6, 138_ = 14.74, *p* < 0.001). Post-hoc tests showed that all regions except aMFG showed increased activation for reward trials compared to no-reward trials (Two tailed, Bonferroni corrected for 7 comparisons: *t_23_* > 3.36, *p* < 0.019, *d_z_* > 0.69 for all ROIs except aMFG; a trend for aMFG: *t_23_* = 2.9, *p* = 0.057, *d_z_* = 0.59). Overall, the MD network showed a strong univariate reward effect during the cue epoch.

We next asked whether the cues were decodable as measured using MVPA, and whether decoding levels increased with reward as has been previously reported (Etzel et al., 2016; Hall-McMaster et al., 2019). Decoding between the two cues separately for the two reward levels were computed in each of the MD ROIs. A two-way repeated measures ANOVA with reward (2) and ROI (7) as factors showed no main effects or interactions (*F* < 1.9, *p* > 0.08). Decoding levels averaged across all MD ROIs were (mean ± SD) 51.71% ± 4.39% and 50.55% ± 6.69% for the reward and no-reward conditions, respectively. There was a marginally significant cue decoding for reward conditions, and no significant decoding for no-reward conditions. (One-tailed t-test, reward: *t_23_* = 1.9, *p* = 0.07, *d_z_* = 0.39; no-reward: *t_23_* = 0.4, *p* = 0.7, *d_z_* = 0.07). Overall, our results show that despite substantial increases in overall univariate activity with reward during the cue epoch across the MD network, the cues were not decodable in both no-reward and reward conditions.

### 3.3 Univariate activity in the MD network during the stimulus epoch

We started our analysis for the stimulus epoch by testing for the effect of reward motivation on the overall activity in MD regions, and whether such effect is different for the three behavioral status conditions. We used averaged β estimates for each behavioral status (Target, High-conflict nontarget, Low-conflict nontarget) and reward level (no-reward, reward) in each of the subject-specific MD ROIs (Figure 3).

**Figure 3:**
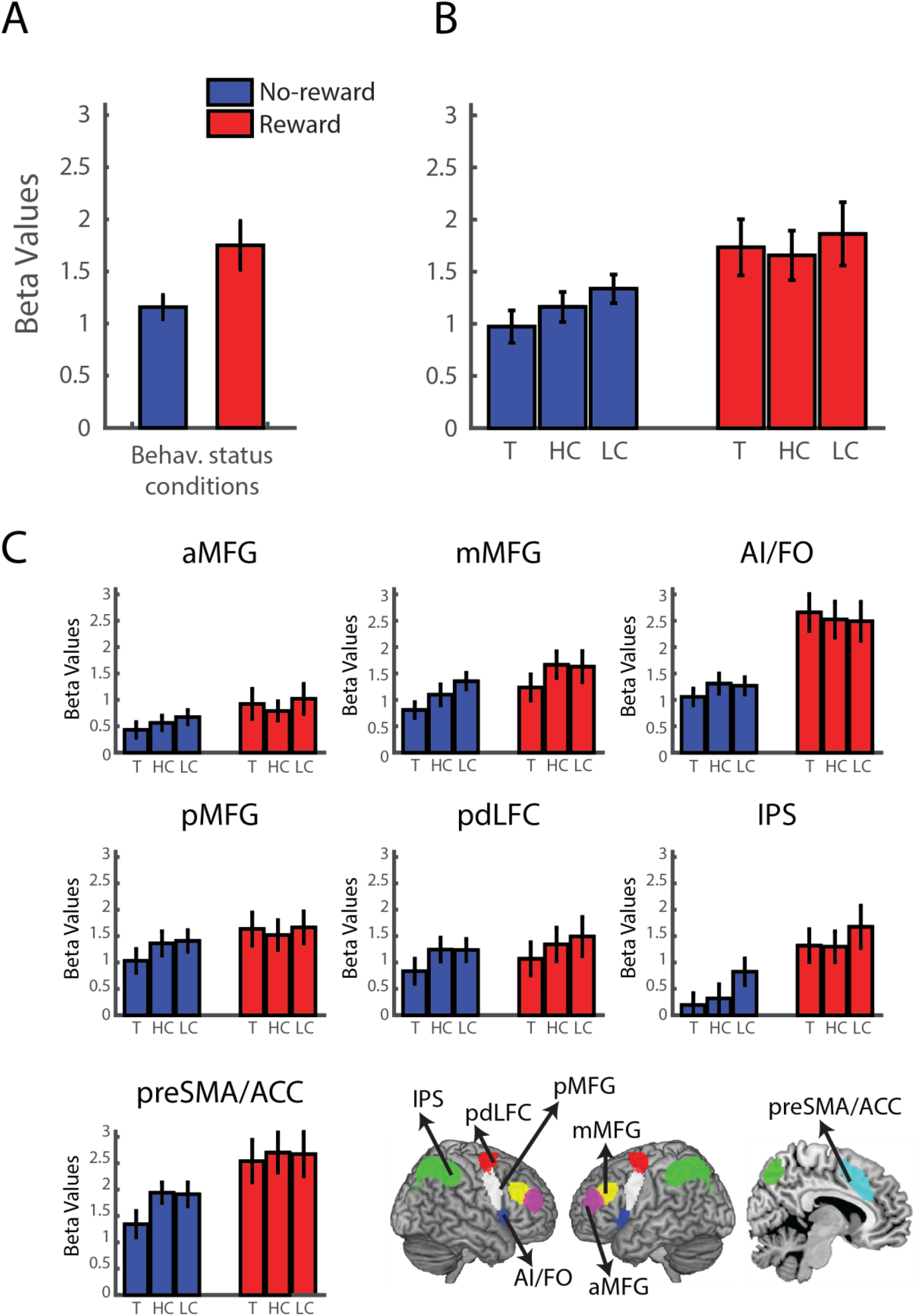
Univariate activity across the MD network during the stimulus epoch. **A.** Univariate results averaged across the subject-specific ROIs for all MD regions. Results are averaged across the behavioral status conditions for no-reward (blue bar) and reward (red bar) conditions. **B.** Average univariate activity across the MD network is shown separately for each behavioral status condition for no-reward (blue bars) and reward (red bars) conditions. T: Target, HC: High-conflict nontarget, LC: Low-conflict nontarget). **C.** Univariate results for the individual MD regions. Post-hoc tests showed that activity increased with reward in the AI/FO, IPS and preSMA, as well as an increase in mMFG that did not survive correction for multiple comparisons. The MD network template is shown for reference. pdLFC: posterior/dorsal lateral prefrontal cortex, IPS: intraparietal sulcus, preSMA: pre-supplementary motor area, ACC: anterior cingulate cortex, AI: anterior insula, FO: frontal operculum, aMFG, mMFG, pMFG: anterior, middle and posterior middle frontal gyrus, respectively. Error bars indicate S.E.M.

Average β estimates across MD regions for both no-reward and reward conditions across all behavioral status conditions were significantly above 0 (Figure 3A), confirming their recruitment during the task, and providing an indication that it was cognitively demanding (no-reward: 1.06 ± 0.58, *t_23_* = 8.85, corrected *p* < 0.001, *d_z_* = 1.81; reward: 1.71 ± 1.22, *t_23_* = 6.83, corrected *p* < 0.001, *d_z_* = 1.39; two-tailed t-test, Bonferroni corrected for 2 comparisons). A three-way repeated measures ANOVA with reward (2), behavioral status (3) and ROI (7) as within-subject factors showed a significant main effect of reward (*F*_1, 23_ = 8.66, *p* = 0.007). There was an interaction of reward level and ROI (*F*_6, 138_ = 14.83, *p* < 0.001), with AI/FO, IPS, and preSMA showing reward effect following post-hoc tests and Bonferroni correction for multiple (7) comparisons (AI: *t_23_* = 5.26, *p* < 0.001, *d_z_* = 1.07; IPS: *t_23_* = 3.31, *p* = 0.021, *d_z_* = 0.68; preSMA: *t_23_* = 3.36, *p* = 0.019, *d_z_* = 0.69). The mMFG showed a reward effect that did not survive multiple comparisons (*t_23_* = 2.26, uncorrected *p* = 0.037, corrected *p* = 0.26, *d_z_* = 0.46). Importantly, there was no main effect of behavioral status (*F*_2, 46_ = 2.25, *p* = 0.12) and no interaction of reward and behavioral status (*F*_2, 46_ = 0.52, *p* = 0.6). Overall, the univariate results indicated increased BOLD response with reward across large parts of the MD network with similar levels of activity for the three behavioral status conditions.

### 3.4 Effect of reward motivation on discrimination of task-related behavioral status in the MD network

Our main question concerned the representation of task-related behavioral status information across the MD network and its modulation by reward, and we used MVPA to address that. For each participant and subject-specific ROI, we computed the classification accuracy above chance (50%) for the distinctions between Target vs. High-conflict nontarget, Target vs. Low-conflict nontarget and High-conflict vs. Low-conflict nontargets, separately for no-reward and reward conditions (Figure 4). The analysis was set to test for discrimination between behavioral status conditions within each reward level, and whether these discriminations are larger when reward is introduced compared to the no-reward condition. A three-way repeated-measures ANOVA with reward (2), behavioral distinction (3) and ROI (7) as within-subject factors showed no main effect of ROI (*F*_6, 138_ = 0.97, *p* = 0.45) or any interaction of ROI with reward and behavioral distinction (*F* < 1.16, *p* > 0.31). Therefore, the classification accuracies were averaged across ROIs for further analysis (Figure 4A). First, we looked at the overall discrimination of behavioral status pairs. Averaged across the three pairs of behavioral status, decoding accuracies were (mean ± SD) 51.4% ± 2.8% and 51.8% ± 3.5% for the no-reward and reward conditions, respectively. Decoding levels were above chance (50%) for both the no-reward and reward trials (one-tailed t-test against chance, corrected for 2 comparisons, no-reward: *t_23_* = 2.34, corrected *p* = 0.03, *d_z_* = 0.48; reward: *t_23_* = 2.5, corrected *p* = 0.02, *d_z_* = 0.5). The decoding levels above chance for the individual pairs of behavioral status for the no-reward and reward conditions are summarized in Table 1. Overall, our results show that on average behavioral status distinctions are represented across the MD network in both no-reward and reward conditions, with some differences between individual pairs of behavioral status.

**Figure 4:**
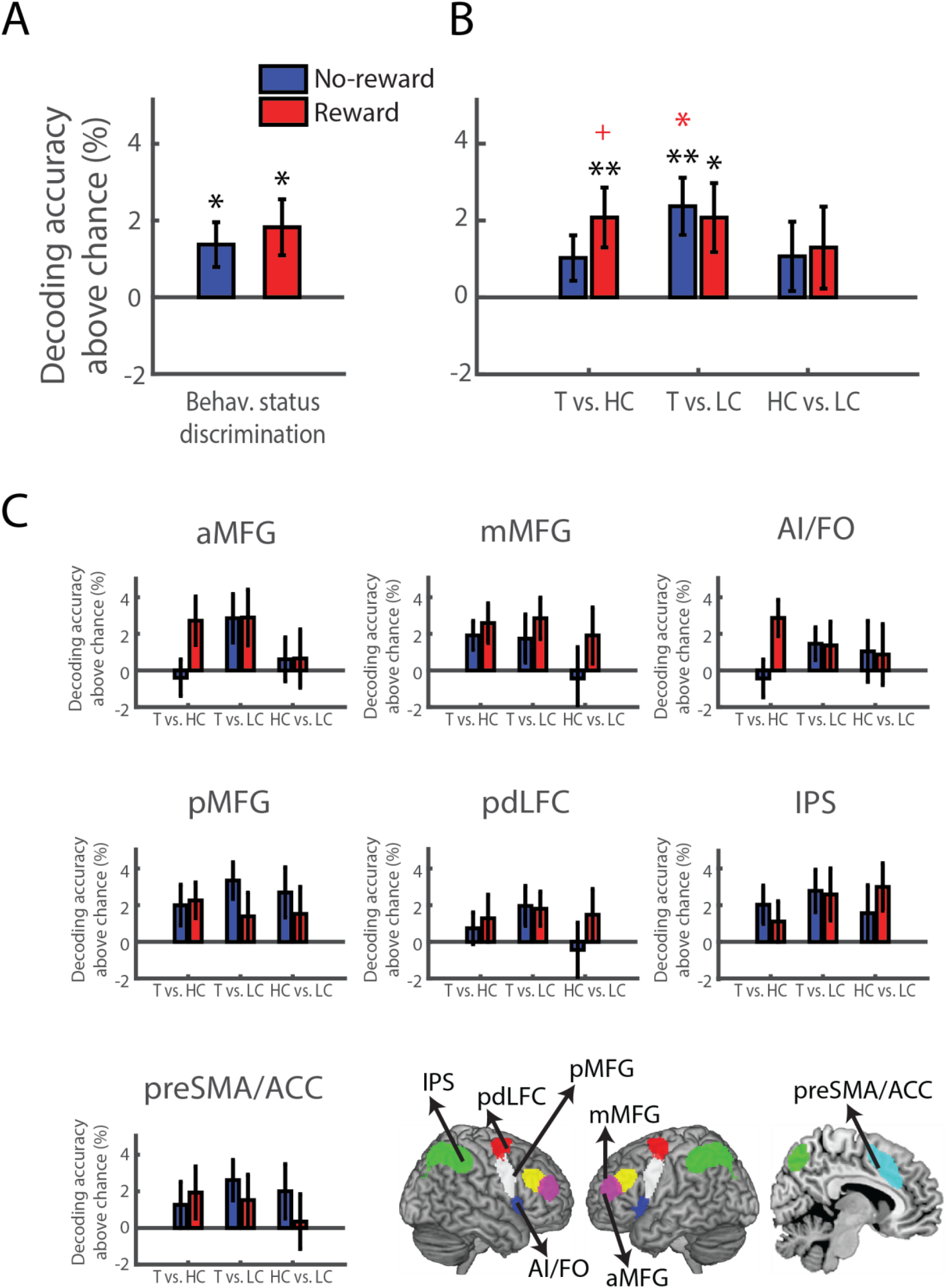
Reward does not modulate distinctions of behavioral status across the MD network. **A.** Classification accuracy is presented as percentage above chance (50%), averaged across all MD regions and behavioral status pairs, for no-reward (blue bars) and reward (red bars) trials. Behavioral status was decodable but not modulated by reward. Asterisks above bars show significant decoding above chance (One-tailed, Bonferroni corrected for 2 comparisons). **B.** The data in A is shown separately for the three distinctions of Target vs. High-conflict nontarget, Target vs. Low-conflict non-target, and High- vs. Low-conflict nontargets. T: Target, HC: High-conflict nontarget, LC: Low-conflict nontarget. Asterisks above bars show one-tailed significant discrimination between behavioral categories above chance without correction (black), and Bonferroni corrected for multiple (6) comparisons (red). See Table 1 for details. **C.** Decoding results are shown for the individual MD regions. The MD network template is shown for reference. pdLFC: posterior/dorsal lateral prefrontal cortex, IPS: intraparietal sulcus, preSMA: pre-supplementary motor area, ACC: anterior cingulate cortex, AI: anterior insula, FO: frontal operculum, aMFG, mMFG, pMFG: anterior, middle and posterior middle frontal gyrus, respectively. Error bars indicate S.E.M. + *p* < 0.06, * *p* < 0.05, ** *p* < 0.01.

**Table 1:**
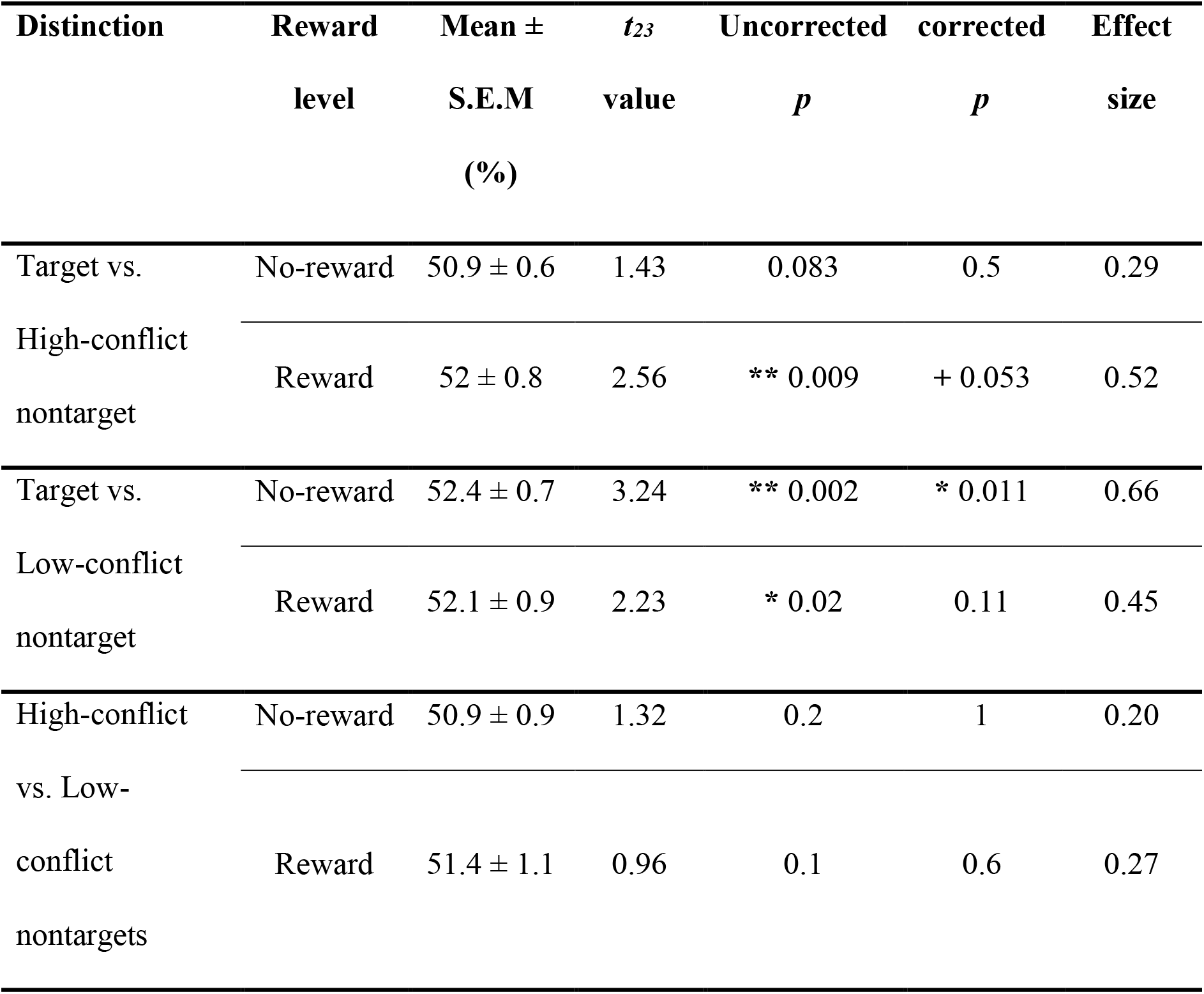
Decoding accuracies for pairs of behavioral status across the MD network. *t* values are for a one-tailed t-test against chance level (50%). Corrected *p* values were obtained using Bonferroni correction for 6 comparisons. + *p* < 0.06, * *p* < 0.05, ** *p* < 0.01

In our critical analysis we tested for the modulatory effect of reward on the discriminability between pairs of behavioral status. In contrast to our prediction, a two-way repeated measures ANOVA with reward (2) and behavioral distinction (3) as within-subject factors showed no main effects of reward or behavioral distinction (*F*_1, 23_ = 0.26, *p* = 0.6; *F*_2, 46_ = 1.37, *p* = 0.26, respectively), and no interaction of the two (*F_2, 46_* = 0.74, *p* = 0.48). In contrast to our prediction, decoding levels across behavioral status pairs were not larger for reward trials compared to no-reward trials (one-tailed paired t-test: *t_23_* = 0.52, *p* = 0.3, *d_z_* = 0.44). To test for the specific prediction that reward might increase discrimination for the high conflict pair of conditions that may not have been picked up by the ANOVA, we compared decoding levels for the Target vs. High-conflict nontarget for the no-reward and reward conditions. Classification accuracy was not larger in the reward trials compared to the no-reward trials for the Target vs. High-conflict nontarget distinction (One-tailed paired t-test: *t_23_* =1.07, *p* = 0.15, *d_z_* = 0.22). A post-hoc sensitivity power analysis yielded an effect size of *d_z_* = 0.52 as the minimum detectable effect. In summary, although the average decoding levels of behavioral status were above chance across the MD network, we did not find increases in decodability with reward. The study had sufficient power to detect a reward modulation with a moderate effect size of 0.52 or more, but not a smaller one.

We conducted several control analyses to confirm the robustness of the results across analysis choices. First, a similar pattern of results was evident across a range of ROI sizes (100, 150, 250 and 300 voxels), confirming that the results do not depend on the choice of ROI size. A two-way repeated measures ANOVA with reward (2) and behavioral distinction (3) as within-subject factors showed no main effects of reward or behavioral distinction and no interaction of the two in any of the ROI sizes (*F* < 1.5, *p* > 0.23). Additionally, the classification accuracy of Targets vs. High-conflict nontargets was not larger in reward trials compared to no-reward trials in any of the ROI sizes (one-tailed paired t-test: *t_23_* < 1.17, *p* > 0.13, *d_z_* < 0.24).

Second, to demonstrate that the results do not depend on our choice to use subject-specific ROIs, we repeated the analysis using the entire MD network template. Overall, the results were similar to those obtained using the subject-specific ROIs. Averaged across all MD ROIs, discrimination of behavioral status was significantly above chance for both no-reward and reward trials (no-reward: 51.6% ± 2.9%, t_23_ = 2.65, corrected *p* = 0.014, *d_z_* = 0.54; reward 52.22% ± 4%, t_23_ = 2.73, corrected *p* = 0.012, *d_z_* = 0.56; one-tailed t-test against chance, Bonferroni corrected for 2 comparisons). A two-way repeated measures ANOVA with reward (2) and behavioral distinction (3) as within-subject factors showed no effect of reward or behavioral distinction (*F*_1, 23_ = 0.44, *p* = 0.51; *F*_2, 46_ = 2.22, *p* = 0.12, respectively), and no interaction of the two (*F*_2, 46_ = 1.21, *p* = 0.31). A specific comparison of classification accuracy for the Target vs. High-conflict nontarget distinction showed that decoding was not larger in the reward trials compared to the no-reward trials (One-tailed paired t-test: *t_23_* =1, *p* = 0.16, *d_z_* = 0.21).

Third, we used another multivariate discriminability measure, linear discriminant contrast (LDC), to assess the difference in dissimilarity of distributed patterns of activity of the behavioral status pairs with and without reward. A larger LDC value for a pair of conditions indicates greater pattern dissimilarity between them, i.e., greater discriminability, with zero indicating no discriminability. Overall, the LDC results were highly similar to decoding results. Behavioral status pairs were significantly discriminable across the MD network in the reward condition and marginally significant after correction in the no-reward condition (averaged across MD ROIs and the three behavioral status pairs, one-tailed t-test against zero, corrected for 2 comparisons, no-reward: *t_23_* = 1.92, corrected *p* = 0.068, *d_z_* = 0.39; reward: *t_23_* = 3.09, corrected *p* =0.005, *d_z_* = 0.63). The discriminability above chance as obtained with LDC (i.e., above zero) for the individual pairs of behavioral status for the no-reward and reward conditions were again highly consistent with the decoding results. Discriminability above chance when uncorrected for multiple comparisons was significantly above chance for Targets vs. High-conflict nontargets and Targets vs. Low-conflict nontargets, for both no-reward and reward conditions (one-tailed t-test: *t_23_* > 1.91, *p* < 0.035), but not for High- vs. Low-conflict nontargets (one-tailed t-test: *t_23_* < 1.5, *p* > 0.075). Discriminations between Targets vs. High-conflict nontargets in the reward condition, and Targets vs. Low-conflict nontargets in the no-reward condition survived Bonferroni correction for 6 comparisons (corrected *p* < 0.016; for the other comparisons: corrected *p* > 0.1).

To test for the effect of reward on pattern discriminability of behavioral status as measured with LDC, we used a two-way repeated measures ANOVA with reward (2) and behavioral distinction (3) as within-subject factors. There were no effects of reward or behavioral distinction (*F*_1, 23_ = 1.03, *p* = 0.32; *F*_2, 46_ = 0.96, *p* = 0.39, respectively), and no interaction of the two (*F*_2, 46_ = 1.36, *p* = 0.27), demonstrating that discriminability was not different between the reward and the no-reward conditions. A further specific comparison of discriminability between Targets and High-conflict nontargets showed a trend of larger pattern dissimilarity for the reward compared to the no-reward condition (one-tailed paired t-test: *t_23_* =1.59, *p* = 0.06, *d_z_* = 0.33).

Fourth, we conducted a complementary whole-brain searchlight analysis to test for increased discriminability of behavioral status with reward across the brain. In a second-level random-effects analysis of behavioral status classification maps (average across the three pairs of behavioral status) of reward vs. no-reward conditions, none of the voxels survived an FDR threshold of p < 0.05. A separate searchlight analysis for classification of Targets vs. High-conflict nontargets showed similar results, with no voxels surviving FDR correction (p < 0.05). Therefore, this analysis did not reveal any other brain regions that showed increase in discriminability with reward.

Fifth, to ensure that our results do not depend on accounting for RT in the GLM regressors, we repeated the analysis using regressors for the stimulus phase with fixed duration (delta function). All the other GLM regressors remained as before. A two-way repeated measures ANOVA with reward (2) and behavioral distinction (3) as within-subject factors showed no main effects of reward or behavioral distinction (*F*_1, 23_ = 0.04, *p* = 0.84; *F*_2, 46_ = 0.85, *p* = 0.44, respectively), and no interaction of the two (*F_2, 46_* = 0.68, *p* = 0.51). There was no increase in classification accuracy in the specific comparison of the Target vs. High-conflict nontarget distinction in reward trials compared to the no-reward trials (One-tailed paired t- test: *t_23_* =0.6, *p* = 0.28, *d_z_* = 0.12).

Lastly, as a quality check for the data and analysis pipeline, we classified right vs. button presses in two motor regions, the precentral gyrus and the supplementary motor cortex. Decoding accuracies averaged across no-reward and reward conditions were high and significantly above chance (50%) in both regions, as would be expected, thus providing support for the good quality of the data and the analysis procedure. (precentral gyrus: 75.35% ± 9.65%, t_23_ = 12.88, corrected *p* = 0.001, *d_z_* = 2.63; supplementary motor cortex: 81.6% ± 12.88%, t_23_ = 12.02, corrected *p* = 0.001, *d_z_* = 2.45; one-tailed t-test against chance, Bonferroni corrected for 2 comparisons).

### 3.5 Effects of reward motivation on task-related behavioral status in LOC

It is widely accepted that the frontoparietal MD network exerts top-down control on visual areas, contributing to task-dependent processing of information. As a comparison to the MD network, we performed similar univariate and multivariate analyses during the stimulus epoch in the high-level general-object visual region, the lateral occipital complex (LOC), separately for its two sub-regions, LO and pFs. We report decoding in subject-specific ROIs driven by the visual categories themselves, as would be expected in LOC, in the following sections.

We first conducted univariate analysis to test for an effect of reward and behavioral status on overall activity in LOC. A four-way repeated-measures ANOVA with reward (2), behavioural status (3), ROI (2), and hemisphere (2) as within-subject factors showed no main effect of reward (*F*_1, 23_ = 2,15 *p* = 0.16), a main effect of ROI (*F*_1, 23_ = 14.70 *p* < 0.001), and a main effect of hemisphere (*F*_1, 23_ = 14.97, *p*< 0.001). There was an interaction of reward and ROI (*F*_1, 23_ = 12.5, *p* = 0.002), and post-hoc tests with correction for multiple (2) comparisons showed that activity was larger for reward compared to no-reward trials in LO (Two-tailed t-test: *t_23_* = 2.55, *p* = 0.036, *d_z_* = 0.52), but not in pFs (Two-tailed t-test: *t_23_* = 0.028, *p* > 0.9, *d_z_* = 0.006). There was a main effect of behavioral status (*F*_2, 46_ = 6.92, *p* = 0.002), but importantly there was no interaction of behavioral status and reward (*F*_2, 46_ = 0.73, *p* = 0.50). Altogether, the univariate results show that reward effects in LOC were partial and limited to LO.

We then tested for the representation of the behavioral status conditions in LOC (Figure 5). Decoding levels averaged across all pairs of behavioral status and the two LOC ROIs were above chance for both no-reward and reward conditions (mean ± SD: 53.23% ± 3.64% and 53.76% ± 4.14% for the no-reward and reward conditions, respectively. One-tailed t-test against chance, corrected for 2 comparisons, no-reward: *t_23_* = 4.34, corrected *p* < 0.001, *d_z_* = 0.9; reward: *t_23_* = 4.45, corrected *p* < 0.001, *d_z_* = 0.9). Importantly, decoding levels were not larger for the reward conditions compared to the no-reward conditions for any of the behavioral status distinctions, with similar results for both LO and pFs. A four-way repeated-measures ANOVA with reward (2), behavioral distinction (3), ROIs (2) and hemispheres (2) as within-subject factors showed no main effect of reward (*F*_1, 23_ = 0.34, *p* = 0.56) or interaction of reward and ROI (*F*_1, 23_ = 1.14, *p* = 0.29). No other main effects or interactions were significant (*F* < 3.15, *p* > 0.05). Overall, these results demonstrate that reward did not modulate the coding of the task-related behavioral status distinctions in LOC.

**Figure 5:**
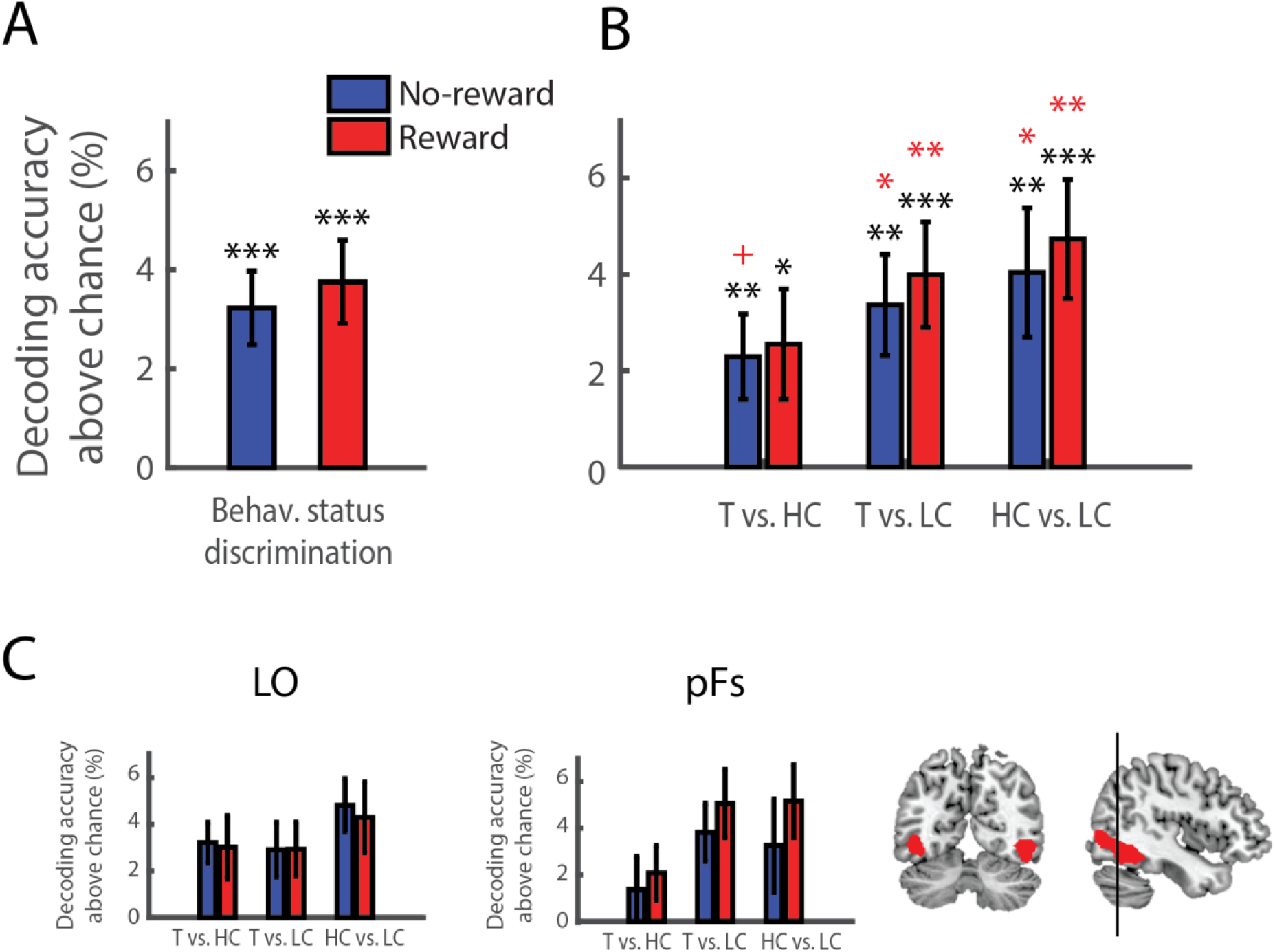
Reward motivation does not increase coding of behavioral status in LOC. **A.** Classification accuracy averaged across all behavioral status pairs is presented as percentage above chance (50%), averaged across LO and pFS and both hemispheres. Classification accuracies for no-reward (blue bars) and reward (red bars) conditions are similar and above chance. Asterisks above bars show significant decoding level above chance, (one-tailed Bonferroni corrected for 2 comparisons). B. Classification accuracies are similar for all three behavioral status distinctions. T: Target, HC: High-conflict nontarget, LC: Low-conflict nontarget. Asterisks above bars show one-tailed significant discrimination between behavioral categories above chance without correction (black), and corrected for multiple (6) comparisons (red). **C.** Classification accuracies for LO and pFs are presented separately, averaged across hemispheres. The LOC template is shown on sagittal and coronal planes, with a vertical line dividing it into posterior (LO) and anterior (pFs) regions. Error bars indicate S.E.M. + *p* < 0.06, * *p* < 0.05, ** *p* < 0.01, *** *p* < 0.001.

### 3.6 Conflict-contingent vs. visual category effects

An important aspect of the Target and High-conflict nontarget conditions in this experiment was that they both contained the same visual categories, which could be either a target or a nontarget (Figure 1B). Therefore, the Target vs. High-conflict nontarget pairs of conditions in our decoding analysis included cases where the stimuli in the two conditions were items from different visual categories (e.g. shoe and sofa following a ‘shoe’ cue), as well as cases where the two stimuli were items from the same visual category (e.g. shoe following a ‘shoe’ cue and a ‘sofa’ cue). We further investigated whether the representation in the MD network and in the LOC was driven by the task-related high conflict nature of the two conditions or by the different visual categories of the stimuli, and whether there was a facilitative effect of reward which is limited to the representation of the visual categories. Figure 6A shows Target vs. High-conflict nontarget distinctions for no-reward and reward conditions, presented separately for pairs of conditions in which the stimuli belonged to the same visual category (different cue trials), and for pairs in which the stimuli belonged to different visual categories (same cue trials), for both the MD and LOC. The data in the figure is the same as in Figure 5 for LOC. For MD, we repeated the same analysis as shown in Figure 4, except for selecting 180 voxels in each subject-specific ROI, in order to keep the ROI size the same for MD and LOC. For both MD and LOC regions, there was no interaction with ROI or hemisphere, therefore accuracy levels were averaged across hemispheres and ROIs for the MD network and LOC (repeated measures ANOVA with reward (2), distinction type (2, same or different visual category), ROIs (7 for MD, 2 for LOC) and hemispheres (2, just for LOC) as within-subject factors: *F* < 3.15, *p* > 0.05 for all interactions with ROI and hemisphere).

**Figure 6:**
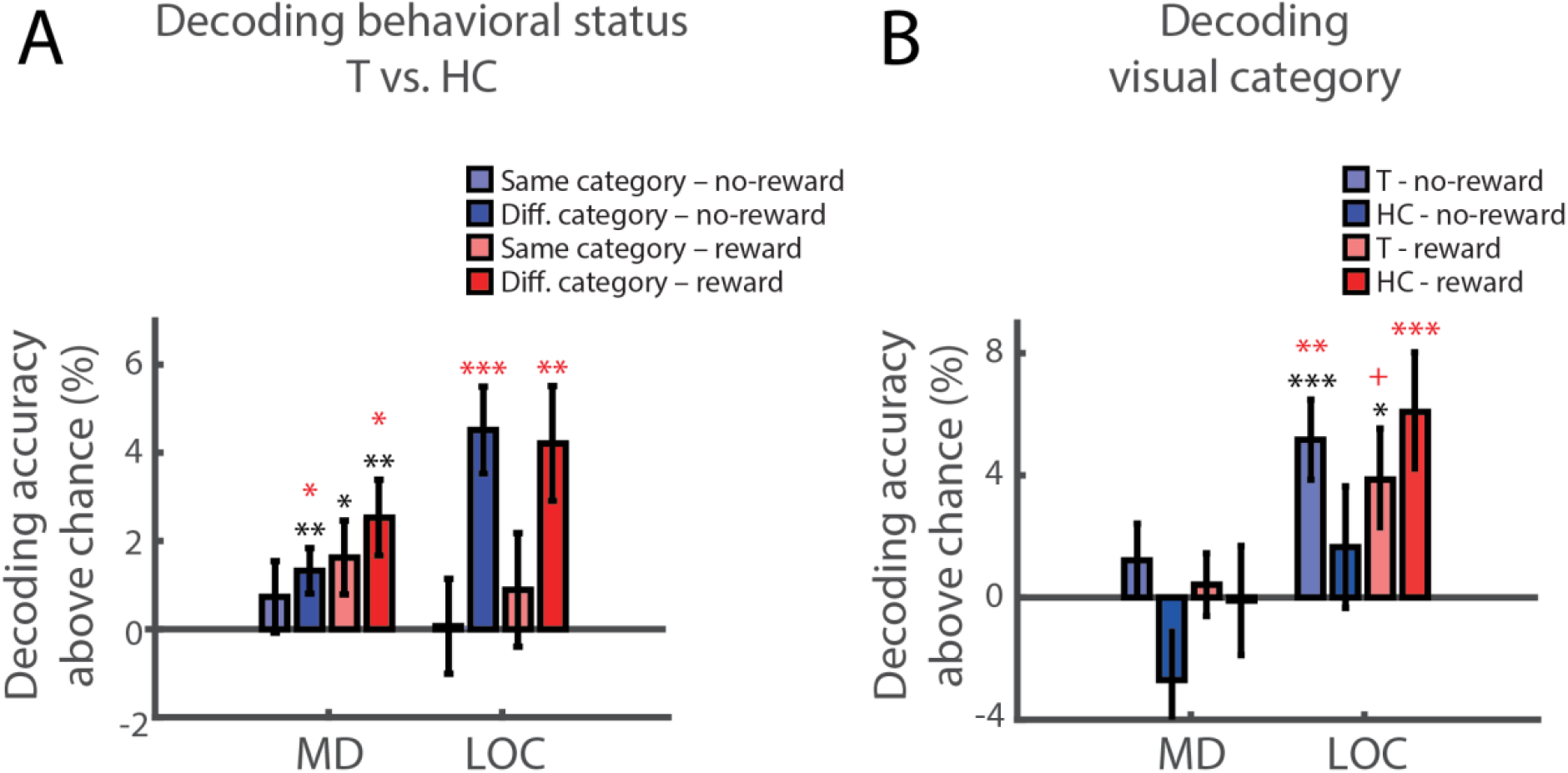
Decoding of visual categories. **A.** Decoding of highly conflicting behavioral status distinctions in the MD network and LOC. Classification accuracies above chance (50%) are presented for no-reward and same-visual-category distinctions (light blue), no-reward and different-visual-category distinctions (dark blue), reward and same-visual-category distinctions (light red), and reward and different-visual-category distinctions (dark red), separately for the MD network and the LOC, averaged across regions and hemispheres in each system. In the MD network, neither reward nor visual category modulated the discrimination of Target vs. High-conflict nontarget. In contrast, classification accuracies in the LOC are larger when the displayed objects are from two different visual categories compared to when they belong to the same visual category, irrespective of the reward level. **B.** Decoding of visual categories with the same behavioral status. Classification accuracies above chance (50%) are presented for no-reward Target categories (light blue), no-reward High-conflict nontargets (dark blue), reward Targets (light red), and reward High-conflict nontargets (dark red), separately for the MD network and for the LOC, averaged across regions and hemispheres in each system. Asterisks above bars show one-tailed significant discrimination above chance without correction (black), and corrected for multiple (4) comparisons (red). Error bars indicate S.E.M. + *p* < 0.06, * *p* < 0.05, ** *p* < 0.01, *** *p* < 0.001.

We next tested for the effect of reward and distinction type (same or different visual category) on decoding levels in each the two systems. In the MD network, a two-way repeated measures ANOVA with reward (2) and distinction type (2) as factors showed no main effect of reward (*F*_1, 23_ = 0.92, *p* = 0.35) and no effect of category distinction or their interaction (*F*_1, 23_ = 2.9, *p* = 0.1; *F*_1, 23_ = 0.1, *p* = 0.8, respectively). These results show that there was no effect of reward on high conflict items that may be specific for the distinction between visual categories. In contrast, a similar ANOVA for LOC showed a main effect of distinction type (*F*_1, 23_ = 25.9, *p* < 0.001) and no effect of reward or their interaction (*F*_1, 23_ = 0.05, *p* = 0.8; *F*_1, 23_ = 0.46, *p* = 0.50, respectively). Together, these results demonstrate that representation was driven by visual categories in LOC, but not in the MD network. To further establish this dissociation between the two systems, we used a three-way repeated measures ANOVA with distinction type (2, same or different visual category), reward (2), and brain system (2, MD or LOC) as within-subject factors. There was no main effect of brain system (*F*_1, 23_ = 1.2, *p* = 0.3), allowing us to compare between the two systems. An interaction between distinction type and system (*F*_1, 23_ = 16.7, *p* < 0.001) confirmed that decoding levels in the two systems were affected differently by visual category. Critically, reward did not lead to increased decoding in either of the systems (no main effect of reward or interactions with reward: *F* < 0.77, *p* > 0.39).

### 3.7 Representation of visual categories

While the main focus of the study was the representation of behavioral status, we further tested whether information about the visual categories themselves is represented across the MD network and in LOC and whether this representation is modulated by reward. Figure 6B shows accuracy levels of visual categories when both have the same behavioral status, i.e. both Targets or both High-conflict nontargets, for no-reward and reward conditions, for both MD and LOC, averaged across ROIs and hemispheres. In the MD network, repeated measures ANOVA with reward (2), behavioral status (Targets, High-conflict nontargets) and ROIs (7) as within-subject factors showed no main effect of reward (*F*_1,23_ = 0.25, *p* = 0.61) or behavioral status (*F*_1,23_ = 2.38, *p* = 0.14). There was a marginally significant interaction of behavioral status and ROI (*F*_6,138_ = 2.17, *p* = 0.049), and post-hoc tests revealed greater decoding for Targets compared to High-conflict nontargets in the preSMA only (corrected *p* = 0.025, corrected for 7 comparisons). Averaged across ROIs, decoding of visual category in the MD network was not above chance when the stimuli were either Targets or High-conflict nontargets, and for either no-reward or reward conditions (*t_23_* < 1.73, corrected *p* > 0.19). These results show that visual categories with the same behavioral status were not represented across the MD network, and this representation did not increase with reward.

In LOC, classification accuracies of visual categories averaged across ROIs and hemispheres was above chance for Targets in the no-reward condition (one-tailed t-test against chance, corrected for multiple (4) comparisons: *t_23_* = 3.94, corrected *p* = 0.001) and marginally above chance in the reward condition (*t_23_* = 2.32, corrected *p* = 0.06). For High-conflict nontargets, decoding was above chance in the reward condition (t*_23_* =3.11, corrected *p* = 0.01) but not in the no-reward condition (*t_23_* = 0.83, corrected *p* = 0.83). Repeated measures ANOVA with reward (2), behavioral status (Targets, High-conflict nontargets), ROIs (2) and hemispheres (2) as within-subject factors showed no main effects or interactions (*F* < 3.94, *p* > 0.059), except for an interaction of behavioral status and reward (*F*_1.23_ = 5.4, *p* = 0.03). However, none of the post-hoc comparisons were significant (corrected *p* > 0.08, corrected for 2 comparisons). Overall, in contrast to the MD network, visual categories in LOC were decodable, at least in part. A further ANOVA demonstrated this dissociation between the systems, with larger accuracy levels in LOC compared to the MD network (repeated measures ANOVA with behavioral status (2), reward level (2) and brain system (2, MD or LOC) as within subject factors: main effect of system: *F*_1.23_ = 18.27, *p* < 0.001; no other significant main effects or interactions: *F* < 3.49, *p* > 0.075). Importantly, despite the larger decodability of visual categories in LOC, there was no effect of reward in any of the systems.

## 4. Discussion

In this study we used a cued target detection task to test for the effect of reward motivation on the coding of task-relevant information in the frontoparietal MD network as reflected in distributed patterns of fMRI data. Reward motivation, in the form of monetary reward, led to overall increase in activity across large parts of the MD network. Using MVPA, we showed that information about the behavioral status during the stimulus epoch of a trial was represented across the MD network. However, in contrast to our prediction, reward motivation did not enhance the distinctions between the three behavioral status conditions across the MD network. Additionally, we did not find evidence for a selective facilitative effect of reward on discriminability of highly conflicting items (competition-contingent effect). In the LOC, information about the behavioral status of the presented stimuli was primarily driven by visual categories, as expected, and was not modulated by reward motivation.

Previous reports showed an enhancement effect of reward on overall activity in the frontoparietal control network (Botvinick and Braver, 2015; Dixon and Christoff, 2012; Padmala and Pessoa, 2011), in line with our data that showed increase in univariate activity with reward during both the cue and stimulus epochs. Our results replicated our previously reported findings that demonstrated the representation of behavioral status information across the frontoparietal cortex (Erez and Duncan, 2015), as measured by decoding distinctions between the behavioral status levels. However, despite increases in overall univariate activity with reward across the MD network, our results did not show an increase in representation in reward trials, and we did not observe a selective increase in representation for the highly conflicting items, namely Targets vs. High-conflict nontargets. Recently, Hall-McMaster et al., (2019) showed some increases in task-relevant stimulus features information when reward levels were high, using distributed patterns in EEG data. In the task used in the current study, we tested for representation of behavioral status of the presented items, rather than stimulus features. We did not observe changes in representation similar to the ones observed by Hall-McMaster et al. There may be several possible reasons for that, including the type of representations that were tested (features vs. behavioral status), multiple differences in the design that contribute to our ability to detect multivariate representations, low spatial specificity in the EEG data compared to the more focused ROIs in our fMRI study, and perhaps the most significant being the limited time window where such differences were observed in EEG that cannot be detected with the low temporal resolution fMRI data.

More generally, several reasons can provide potential explanations for the results obtained in our study, showing no facilitative effect of reward on pattern discriminability of behavioral status. Indeed, it is possible that in contrast to reward effects on overall activity, its effect on neural representations is limited to cue decoding when the task context is set, as has been previously demonstrated (Etzel et al., 2016), and does not extend to the stimulus phase when information is processed based on the cue. Another possibility may be related to the reward being offered on a trial-by-trial basis, which may not be sufficiently strong to generate detectable reward-related effects on decoding. It has been recently demonstrated that stable longer-term cue-reward associations that are learned prior to scanning lead to increased decoding of task-related information in visual and parietal areas (Tankelevitch et al., 2020). Other possible explanations may include a reward manipulation that was not sufficiently strong to make a difference to pattern discriminability, and multiple factors in the experimental design that make small effects hard to detect with current MVPA methods, particularly across the frontoparietal cortex where low decoding accuracies are widely common for reasons that are not yet well understood (Bhandari et al., 2018). We note, however, that the results were consistent across several control analyses, including ROI size, multivariate discriminability measure, and whole-brain searchlight analysis. Beyond the above potential explanations, insufficient power is always a concern when reporting null results. A post-hoc sensitivity power analysis showed that the study had sufficient power to detect a moderate effect size with respect to reward modulation of decoding behavioral distinctions, but not a smaller one. For fMRI MVPA results in general, and more so given the sparse evidence in the literature relevant for our main research question, it is hard to establish an expected effect size. Therefore, a reasonable interpretation of our results would be that reward does not lead to moderate or large increase in decoding but could possibly lead to increased decoding if this is relatively small. We note that the study had sufficient power to detect univariate effects as well as other complementary pattern discriminations. First, our overall decoding levels for behavioral status were above chance, with decoding for the no-reward conditions similar to previously reported results (Erez and Duncan, 2015). Second, decoding in LOC showed a clear pattern of contingency on visual category representation, as expected in the visual cortex, in contrast to distinct pattern of decoding across the MD network that did not depend on the visual category of the presented items. Such decoding patterns in LOC are consistent with previous studies that showed only weak, or non-existing, task effects (Bugatus et al., 2017; Harel et al., 2014; Hebart et al., 2018). Lastly, in a control analysis we observed high decoding accuracies for motor responses (right vs. left button presses) in the motor cortex, as expected for these regions. This provides an indication for the quality of the data and the analysis procedure and demonstrates sufficient power to detect these responses.

Our predictions were based on the sharpening and prioritization account, which postulates that reward motivation leads to a sharpened neural representation of relevant information depending on the current task and needs. Previous neurophysiological evidence provide support for this aspect: reward has been associated with firing of dopaminergic neurons (Bayer and Glimcher, 2005; Schultz et al., 1997), and dopamine has been shown to modulate tuning of prefrontal neurons and to sharpen their representations (Ott and Nieder, 2016; Thurley et al., 2008; Vijayraghavan et al., 2007). The prioritization aspect can be related to the expected value of control (EVC) theory (Shenhav et al., 2013) and reward-based models for the interaction of reward and cognitive control, essentially a cost-benefit trade-off (Botvinick and Braver, 2015). Cognitive control is effortful and hence an ideal system would allocate it efficiently, with a general aim of maximizing expected utility. Despite the appeal of this account, our results did not show experimental support for this view. At the behavioral level though, we observed some evidence for such a benefit of reward. Accuracy levels of performance in the task for Target trials were higher in the reward compared to the no-reward condition. Additionally, while in the no-reward condition Target trials were less accurate than Low-conflict nontargets, as is well established in the visual search literature (Schneider and Fisk, 1983; Shiffrin and Schneider, 1977), there were no differences between them in the reward condition. We did not observe a similar benefit in reaction times, most likely due to the time threshold that we used for reward trials, which reduced the reaction time on all the reward trials and may have masked an interaction with reward.

The nature of the conflict addressed in our study concerns the effect of current task goals (i.e., cue) on the behavioral relevance and processing of the presented stimuli. This is a fundamental aspect of cognitive control which have been addressed by us and others in the field in different ways, both in human and non-human primate studies (Erez and Duncan, 2015; Freedman et al., 2001; Hall-McMaster et al., 2019; Kadohisa et al., 2015). There are several other important aspects of the potential effect of reward motivation on goal-directed information processing that cannot be addressed in our data and may be addressed in future studies. For example, one aspect is related to how reward motivation may bias response towards reward-associated stimuli when stimuli are presented simultaneously and compete for attention. Another aspect is how reward motivation may affect encoding of visual features of the stimulus, e.g., as can be tracked using inverted encoding models (Myers et al., 2015; Sprague et al., 2016). These aspects are complementary and provide different perspectives to better understand control processes.

The visual categorization aspect of our task allowed us to investigate effects of reward on representation in LOC compared to the MD network, and in particular whether there is a specific effect of reward that is driven by visual differences. In the MD network, decoding levels were similar between conditions with the same visual category and different visual category, and there was no modulation by reward in any of them. In contrast, the discrimination in LOC was driven by the visual categories, as expected in the visual cortex, with Targets and High-conflict nontargets being discriminable only when items belonged to two different visual categories. In a complementary analysis, we tested for representation of the visual categories themselves, while keeping the behavioral status fixed, i.e., when both stimuli are Targets or both are High-conflict nontargets. As expected, decoding levels of visual categories were larger in LOC compared to MD. Additionally, visual categories with the same behavioral status were not represented in the MD network, in line with previously reported findings (Erez and Duncan, 2015). Importantly, reward did not lead to increased decodability of the visual categories in either the MD network or in the LOC. While it is widely agreed that the frontoparietal cortex exerts top-down effects on visual areas, there is no clear prediction as to whether any effects of reward should be observed in the visual cortex. Our results provide evidence that the effects of reward were not present in LOC. Previous studies have shown differences in representations between pFs and LO (Harel et al., 2014; Jiang et al., 2007; Li et al., 2007), however, our results were similar for both regions.

Although our primary question addressed the representation of task-related information during the integration of stimulus and cue, we also tested for an effect of reward in the cue epoch. The use of catch trials ensured that the cue and stimulus GLM regressors were appropriately decorrelated. The overall univariate activity across the MD network increased with reward during the cue epoch, possibly reflecting an increase in cognitive effort due to the reward. However, we did not observe cue decoding above chance, in contrast to previously reported results (Etzel et al., 2016). One reason for this difference may be related to the design of the current task. We used words of the category names as cues, which appeared together at the same time with the no-reward/reward indication – the reward trial cues had additional red pound signs. The choice of words as cues allowed for high task performance (compared to using abstract symbols, which is more difficult), as confirmed in pilot experiments. This may have come at the expense of cue decodability, which was not the focus of the study. The visually salient reward signal presented simultaneously with the cues may have also contributed to reduced cue decoding levels, and the longer delay period in their study probably led to longer active maintenance of the task rule in working memory. Other reasons for the different results compared to Etzel et al. may be related to the size of the effect and our ability to detect it. The effect of reward on cue decoding reported by Etzel et al. was observed across all ROIs, but was inconsistent in individual ROIs, with only one region showing a significant difference between reward and no-reward conditions. Even for decoding across all ROIs, statistical significance was reached in one statistical test but was only marginal in another. It could be that the effect of reward on cue decoding is relatively small and therefore hard to detect. Among others, our ability to detect such effect may be affected by the areas chosen, number of trials and runs completed per subject etc. The contributions of many of these factors to decoding levels in the frontoparietal network more generally are not yet well understood (Bhandari et al., 2018). The similarities and differences in task design and results that we report here together with those reported previously (Etzel et al., 2016) may be used to better understand the factors that affect decoding levels of contextual cues across the frontoparietal network and inform future studies. Enhancement of cue decoding following reward was also recently reported using EEG (Hall-McMaster et al., 2019). This facilitative effect was observed for ‘switch’ trials but not ‘stay’ trials. Our study design was not controlled for ‘switch’ and ‘stay’ trials, and it could be that cue decoding would emerge if these could be considered. An additional prominent difference between the EEG results and ours is the spatial specificity, with much more widespread activity contributing to decoding in EEG data.

There is a growing interest of the scientific community in the interaction between reward motivation and control processes, its neural correlates, and its implementation in computational models of reinforcement learning. Here, we report increases in overall univariate activity with reward but primarily null results for an effect of reward on task-related representations across the frontoparietal network and we believe that these results may be useful for future studies that seek to address similar questions. Our findings point out that the extent of effects of reward on neural representation may be limited and dependent on specific task demands and trial epochs. Limited effects were also reported by Hall-McMaster et al., (2019), where increased cue decoding was observed only when the cue changed from one trial to another, but not in trials where the cue remained the same. Reports on the effect of reward on task representation are so far limited, and with the movement towards open and replicable science, it is important to establish the best possible pool of evidence for such effects. Our results will ultimately contribute to the overall estimates of the extent of effects of reward on neural representation. Additionally, they may be used as a starting point for future studies and particularly for power estimates, with respect to both the expected effect size as well as aspects of the experimental design that may contribute to the detectability of reward effects. Different measures can be taken to further increase power within scanning time limits. These might include, but not limited to, collecting a larger amount of data per participant, possibly over more than one scanning session, as well as using other methods that have been used recently to address neural representation for fMRI data such as repetition suppression (Garvert et al., 2015).

## Conclusions

With growing interest in the interaction between control processes and reward motivation, our study provides important experimental evidence for the limited extent of effects of reward on task-relevant neural representations. Future studies will be required to further investigate the factors that determine how and when reward shapes neural representations and the mechanisms that underlie the profound effects of reward on behavior.

## Acknowledgements

This work was supported by a Royal Society Dorothy Hodgkin Research Fellowship (UK) to Yaara Erez (DH130100); a Gates Cambridge Trust (Cambridge, UK) scholarship to Sneha Shashidhara; and the Medical Research Council (UK) (Intramural Program MC-A060-5). We thank John Duncan and Daniel Mitchell for fruitful discussions and advice throughout the study. The funding sources had no involvement in any aspects of the study.

## Abbreviations

MD: multiple-demand
LOC: lateral occipital complex
LDC: linear discriminant contrast

### Additional neuroanatomical abbreviations

pdLFC: posterior/dorsal lateral prefrontal cortex
aMFG: anterior part middle frontal gyrus
mMFG: middle part of the middle frontal gyrus
pMFG: posterior part of the middle frontal gyrus
IPS: intraparietal sulcus
preSMA: pre-supplementary motor area
ACC: anterior cingulate cortex
AI: anterior insula
FO: frontal operculum
pFs: posterior fusiform region
LO: lateral occipital region

